# Endometrial receptivity revisited: endometrial transcriptome adjusted for tissue cellular heterogeneity

**DOI:** 10.1101/357152

**Authors:** Marina Suhorutshenko, Viktorija Kukushkina, Agne Velthut-Meikas, Signe Altmäe, Maire Peters, Reedik Mägi, Kaarel Krjutškov, Mariann Koel, Juan Fco. Martinez-Blanch, Francisco M. Codoner, Felipe Vilella, Carlos Simon, Andres Salumets, Triin Laisk

## Abstract

**STUDY QUESTION:** Does cellular composition of the endometrial biopsy affect the gene expression profile of endometrial whole-tissue samples?

**SUMMARY ANSWER:** The differences in epithelial and stromal cell proportions in endometrial biopsies modify whole-tissue gene expression profiles, and also affect the results of differential expression analysis.

**WHAT IS ALREADY KNOWN:** Each cell type has its unique gene expression profile. The proportions of epithelial and stromal cells vary in endometrial tissue during the menstrual cycle, along with individual and technical variation due to the way and tools used to obtain the tissue biopsy.

**STUDY DESIGN, SIZE, DURATION:** Using cell-population specific transcriptome data and computational deconvolution approach, we estimated the epithelial and stromal cell proportions in whole-tissue biopsies taken during early secretory and mid-secretory phases. The estimated cellular proportions were used as covariates in whole-tissue differential gene expression analysis. Endometrial transcriptomes before and after deconvolution were compared and analysed in biological context.

**PARTICIPANTS/MATERIAL, SETTING, METHODS:** Paired early- and mid-secretory endometrial biopsies were obtained from thirty-five healthy, regularly cycling, fertile volunteers, aged 23 to 36 years, and analysed by RNA sequencing. Differential gene expression analysis was performed using two approaches. In one of them, computational deconvolution was applied as an intermediate step to adjust for epithelial and stromal cells’ proportions in endometrial biopsy. The results were then compared to conventional differential expression analysis.

**MAIN RESULTS AND THE ROLE OF CHANCE:** The estimated average proportions of stromal and epithelial cells in early secretory phase were 65% and 35%, and during mid-secretory phase 46% and 54%, respectively, that correlated well with the results of histological evaluation (r=0.88, p=1.1×10^−6^). Endometrial tissue transcriptomic analysis showed that approximately 26% of transcripts (n=946) differentially expressed in receptive endometrium in cell-type unadjusted analysis also remain differentially expressed after adjustment for biopsy cellular composition. However, the other 74% (n=2,645) become statistically non-significant after adjustment for biopsy cellular composition, underlining the impact of tissue heterogeneity on differential expression analysis. The results suggest new mechanisms involved in endometrial maturation involving genes like *LINC01320, SLC8A1* and *GGTA1P*, described for the first time in context of endometrial receptivity.

**LIMITATIONS, REASONS FOR CAUTION:** Only dominant endometrial cell types were considered in gene expression profile deconvolution; however, other less frequent endometrial cell types also contribute to the whole-tissue gene expression profile.

**WIDER IMPLICATIONS OF THE FINDINGS:** The better understanding of molecular processes during transition from pre-receptive to receptive endometrium serves to improve the effectiveness and personalization of assisted reproduction protocols. Biopsy cellular composition should be taken into account in future endometrial ‘omics’ studies, where tissue heterogeneity could potentially influence the results.

**TRIAL REGISTRATION NO:** N/A

## Introduction

The endometrium is a unique self-renewing tissue that undergoes histologically and functionally distinguished phases of growth and differentiation during the menstrual cycle. This is necessary to synchronize the endometrium with oocyte maturation, fertilization, and embryo implantation, which takes place in the mid-secretory phase of menstrual cycle, corresponding to the window of implantation – WOI (Harper, 1992). To understand and characterize the receptive phenotype of endometrium, initially a histological dating method was developed (Noyes *et al.*, 1975). With ‘omics’ technologies, the understanding of molecular processes behind endometrial receptivity has evolved rapidly (Altmäe *et al.*, 2014). Based on this knowledge, the first gene expression-based clinical endometrial receptivity test was introduced in 2011 (Díaz-Gimeno *et al.*, 2011) and has been successfully applied to detect the most appropriate day for embryo transfer in women undergoing IVF (Ruiz-Alonso *et al.*, 2013). Yet, while previous endometrial transcriptome studies, using either microarray or RNA sequencing (RNA-seq) technology, have identified numerous transcripts differentially expressed during the mid-secretory phase, the overlap between the reported significantly altered genes among studies has remained modest and the exact mechanisms of receptivity are still not fully understood (Gómez *et al.*, 2015).

However, given the phase-dependent changes in tissue cellular composition during the menstrual cycle, the gene expression differences identified in the whole-tissue sample may be partially consequent on these changes eclipsing the true alterations in expression levels. This phenomenon is further exaggerated by inter-sample differences in cellular heterogeneity levels, likely due to biological or technical issues, related to the means used to obtain the tissue biopsy. Therefore, for studies reporting differential expression, it is unclear whether the observed changes in transcript levels reflect variations in cellular subpopulations in the whole tissue biopsy, altered gene expression in these subpopulations, or a combination of the two. In support of this idea, distinct gene expression patterns of two dominant endometrial cell types, e.g. epithelial and stromal cells, during WOI were first shown by a study using laser microdissection microscopy to separate endometrial glandular epithelial and stromal cells (Evans *et al.*, 2012), and confirmed by our recent study using transcriptomics of fluorescence-activated sorted cells (Altmäe *et al.*, 2017). However, in addition to costly and time-consuming cell sorting procedure, cell isolation entails a loss of a more general view on tissue complexity. Moreover, removing the cells from their natural niche may alter their genome activity even if the mildest tissue disaggregation conditions are applied and the sorting time is kept short. Although the abovementioned issues support the whole tissue transcriptome analysis, ignoring the cellular heterogeneity in whole tissue analysis may cause misinterpretation of the genomic data, particularly for highly heterogenic tissues (Shen-Orr and Gaujoux, 2013). As an alternative to cell type isolation techniques, computational decomposition (or deconvolution) of gene expression profiles from heterogeneous tissue has been proposed to adjust for differences stemming from variation in biopsy cellular composition, and has been successfully used for gene expression profiling of blood, lymphoid and tumour tissue (Chen *et al.*, 2017b; Li *et al.*, 2017; Pan *et al.*, 2017; Schelker *et al.*, 2017).

In the current study, we utilized a computational approach to estimate cell-type fractions in biopsies and to deconvolute the endometrial whole-tissue gene expression profile. This approach helps to adjust for expression changes stemming from altered cellular composition of biopsied samples taken at different phases of the menstrual cycle, and to reveal the true gene expression changes irrespective of the cellular heterogeneity of the tissue analysed. The current study is the first using gene expression profiling adjusted for cellular heterogeneity to uncover the mechanisms of endometrial receptivity that otherwise would remain hidden in whole-tissue genomic analysis.

## Materials and Methods

The study was approved by the Research Ethics Committee of the University of Tartu, Estonia (No 221/M-31) and Ethical Clinical Research Committee of IVI Clinic, Valencia, Spain (No 1201-C-094-CS). Informed consent was signed by all women who entered the study.

### Study participants

The endometrial tissue samples were obtained in Estonia and Spain from fertile-aged women with normal body mass index (BMI) and self-reported regular menstrual cycles (Estonia n=20, Spain n=15; a total of 35 women) who had not received hormonal treatments for at least three months prior to the time of biopsy. The average age and BMI were 29.1±3.6 years and 23.2±2.9 kg/m^2^ for Spanish women, and 30.2±3.4 years and 23.2±4.5 kg/m^2^ for Estonian women, respectively. The women had at least one live-born child and no previous infertility records. Two endometrial biopsies were obtained using Pipelle catheter (Laboratoire CCD, France) within the same natural menstrual cycle; the first during the early secretory endometrial phase (ESE; two days after the luteinizing hormone surge [LH+2] in Spain and LH+1 to LH+3 in Estonia) and the second sample during the mid-secretory endometrial phase (MSE; day LH+7 in Spain and day LH+7 to LH+9 in Estonia). Endometrial sample timing was confirmed by histological examination according to Noyes’ criteria (Noyes *et al.*, 1975). The samples were stored at - 80°C until use.

### Total RNA extraction from endometrial tissue

For the total RNA extraction, up to 30 mg of the tissue was homogenized in the presence of QIAzol reagent (Qiagen, Germany) and mechanical lysis (0.5 mm diameter steel bead, Tissue-Lyser LT (20-40s)), and processed using miRNeasy Mini kit (Qiagen), following manufacturer’s protocol. miRNA content was decreased using Qiagen’s MinElute protocol. DNase I treatment was performed on column using RNase-Free DNase Set (Qiagen). Purified RNA integrity number (RIN) and quantity were determined with Bioanalyzer 2100 RNA Nano 6000 kit (Agilent Technologies, USA).

### mRNA sequencing

The endometrial total RNA samples with concentration at least 200 ng/µl and RIN >8.0 were used for cDNA synthesis and library construction. Libraries were generated from 4 µg of total RNA using Illumina TruSeq Stranded technology (Illumina, USA), following manufacturer’s protocol. Libraries were normalized, pooled and sequenced using HiSeq™ 2000/2500 with a configuration of 100 cycles paired-end reads in three sample groups: Estonian Est 1 cohort samples at Estonian Genome Center Core Facility (Tartu, Estonia), Est 2 cohort samples at Institute for Molecular Medicine Finland (FIMM, Helsinki, Finland) and Spanish samples at Lifesequencing S.L. (Valencia, Spain).

### RNA sequencing analysis and data preparation

The processing of raw RNA reads, that included quality control, adapter removing, trimming and mapping to human genome hg19 was performed as previously described (Altmäe et al., 2017). The RNA-seq data presented in this study is deposited in the Gene Expression Omnibus database with accession number GSE98386.

### Transcriptome profiling of early secretory and mid-secretory endometrial samples

To rule out population- and/or batch-specific effects, differential expression (DE) analysis was carried out separately in three groups according to sequencing centre: Est 1, Est 2 and Spain. For group-level DE analyses, the edgeR software was used. Before DE analysis, transcripts with low or uneven expression patterns were filtered out, retaining transcripts with counts per million (CPM) ≥ 2 in at least half of the samples (±5%) for subsequent analysis. We used a paired analysis design, which takes into account the paired nature of the biopsies collected from each woman and thus increases statistical power, while at the same time reduces the effect of potential confounding factors.

Thereafter, results from all three participating groups were meta-analysed using the METAL tool (Willer *et al.*, 2010) to identify transcripts that exhibit similar DE patterns in different groups. Genes with consistent effect direction over three groups with p-value lower than Bonferroni corrected p-value threshold (0.05 divided by the number of transcripts included in the meta-analysis) were considered statistically significantly differentially expressed.

### Deconvolution analysis

For the deconvolution of endometrial whole-tissue gene expression profiles, single-cell tagged reverse transcription (STRT)-based RNA-seq data for bulk RNA from endometrial stromal and epithelial cells was used. STRT data for both Estonian and Spanish samples was used to deconvolute the gene expression profiles of samples from respective groups. Totally, we used pre-analysed RNA-seq data for 19 stromal and 17 epithelial fractions from ESE, and 24 stromal and 24 epithelial fractions from MSE. The detailed information on participants and sampling is presented in our previous study (Altmäe *et al.*, 2017). Briefly, the cell populations were extracted by fluorescence-activated cell sorting (FACS) from endometrial biopsies of healthy women, and analysed using the STRTprep pipeline as previously described (Krjutškov *et al.*, 2016) (Figure 1). As the pipeline uses hg19 reference genome for mapping, and a different version of annotation file for aligning/counting, only the .bam files from the pipeline were used for further steps. The mapped reads were aligned on Ensembl v.75 annotation file, the same file used for whole tissue RNA-seq data, and raw counts were obtained with HTSeq package script htseq-count v.0.6.1 (Anders *et al.*, 2015). The normalized CPMs were obtained using R package edgeR v. 3.6.2 (Robinson *et al.*, 2010) and the default normalization function instead the spike-in normalization.

**Figure 1.**
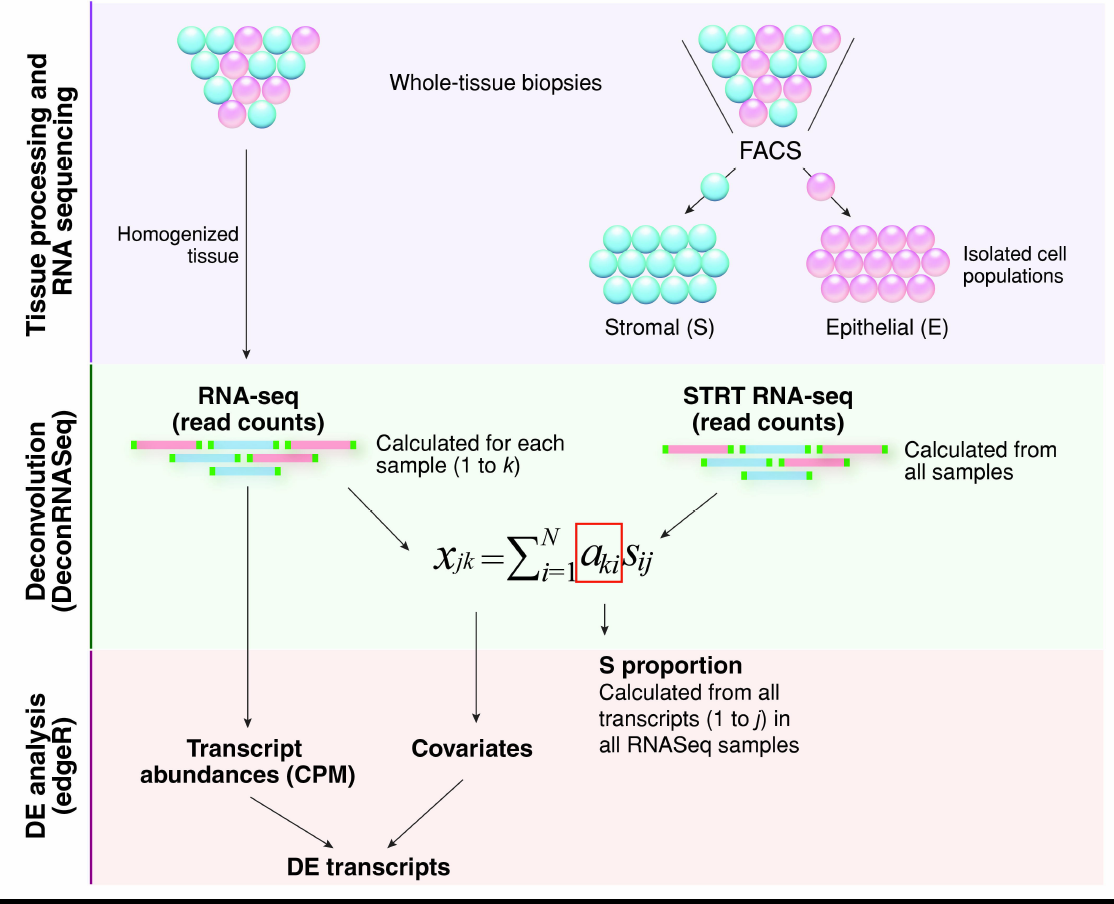
A simplified overview of the gene expression profile computational deconvolution of endometrial tissue. Endometrial biopsies were homogenized and total RNA extracted from whole-tissue samples was sequenced. In parallel, two cell types from endometrial biopsies were separated; epithelial cells were labelled with fluorochrome-conjugated mouse anti-human CD9 monoclonal antibody and stromal cells were simultaneously labelled with fluorochrome-conjugated mouse anti-human CD13 monoclonal antibody, followed by flow cytometric analysis and FACS cell sorting. Bulk-RNA full transcriptome analysis of FACS sorted endometrial epithelial and stromal cells was performed with the RNA-seq method, following the single-cell tagged reverse transcription (STRT) protocol with modifications. Reference proportions of epithelium (E) and stroma (S) were calculated for each sample using DeconRNASeq algorithm. By using transcript measures from separate cell-types (FACS-sorted epithelial and stromal cell populations) and transcript measures from whole-tissue samples, the algorithm estimates the cell-type proportions over samples by fitting a non-negative least squares equation for each transcript. Further, it calculates the theoretical proportions of epithelium and stroma across the whole transcriptome. The expression level *x_jk_* of gene *j* in a sample *k* is the average of expected expression levels across the cell types *s_ij_*, weighted by the respective cell-type proportions *a_ki_* (*i*= 1 … *N*, *N*: the total number of cell types) (Adapted from (Gong and Szustakowski, 2013)). Average stroma proportions calculated from all whole-tissue samples used for RNA-seq were then applied as covariates in differential expression (DE) analysis using edgeR.

For deconvolution, we used the R package DeconRNASeq v.1.10.0 (Gong and Szustakowski, 2013), which uses tissue and pure cell-type specific expression patterns to decompose mixed tissue to cell types. Package adopts a globally optimized non-negative decomposition algorithm through quadratic programming to estimate cell-type proportions in the heterogeneous tissue. As the input it takes normalized expression levels from pure cell fractions for some set of transcripts and the mixed tissue normalized expression levels. Both the cell fraction-specific and mixed expression values should be in same units and normalized using the same method. In this study, normalized CPMs of all non-zero expressed transcripts from stroma and epithelium STRT-sequencing, and from endometrial biopsies were used as the input, and the median value of cell-specific expression per cell-type over sample group was used to estimate the proportions of stromal and epithelial cells in the heterogenous endometrial tissue (Figure 1).

The stromal and epithelial cell proportions obtained from deconvolution for ESE and MSE samples were used in downstream DE analyses. The DE analyses were done the same way as described above, except this time cell proportions were used as covariates (Figure 1). As the stroma and epithelium proportions together add up to 1.0, only the stroma proportions were used as covariates. The meta-analysis for groups was done as described above.

### Correlation between estimated cellular proportions and histological dating

To test the accuracy of the estimated proportions of stromal and epithelial cells resulting from deconvolution, we correlated the epithelial fraction estimates with estimated endometrial cycle day (corresponding to a 28-day cycle) from histological analysis. For this, we used the histological evaluation data for 18 paired samples (nine from ESE and nine from MSE phase) for which detailed dating information was available. If the endometrial cycle date had been given as a range (e.g. days 21-23), the arithmetic mean was used for calculating the Pearson correlation coefficient. Additionally, we assessed the proportion of epithelial and stromal tissue in the two time-points in histological images, and compared the numbers to those obtained from deconvolution analysis. The histological specimens were prepared from formalin-fixed paraffin-embedded endometrial tissue sections using standard hematoxylin and eosin staining protocol (Fischer *et al.*, 2008). The proportions of epithelial and stromal cells in the specimen were robustly calculated by measuring the areas occupied by either cell type in histological images using Adobe Photoshop CC 2017. The size of the selected areas was converted to pixels, and the proportion of epithelial cells was calculated by dividing the area under epithelial cells (in pixels) to that of the entire specimen. The analysis included a total of 13 specimens (eight ESE and five MSE), as for some individuals, more than one image was available for evaluation. In case of multiple measurements for one biopsy, the average proportions were calculated across evaluations.

### Functional annotation of differentially expressed genes

Functional annotation was performed with Ingenuity Pathway Analysis (IPA) tool using all significant differentially expressed genes (DEGs) identified after deconvolution. Pathway analysis was performed using IPA with default settings, and pathways with a Benjamini-Hochberg p-value <0.05 were considered as significantly enriched. Networks of significantly DEGs were then algorithmically generated based on their connectivity.

### Endometrial receptivity markers

To outline potential biomarker candidates that exhibit true expression change in MSE and could thus be detected by routine transcriptome analysis of whole tissue biopsy, meta-analysis DEG lists identified before and after deconvolution analysis (|log_2_FC|≥1, outranked Bonferroni p-value) were compared and overlapping transcripts were selected as marker candidates.

Further on, our marker candidate list was compared with DEG lists reported in previous RNA-seq and microarray studies on human endometrium (Carson *et al.*, 2002; Kao *et al.*, 2002; Borthwick *et al.*, 2003; Riesewijk *et al.*, 2003; Mirkin *et al.*, 2005; Talbi *et al.*, 2006; Altmae *et al.*, 2010; Tseng *et al.*, 2010; Díaz-Gimeno *et al.*, 2011; Hu *et al.*, 2014; Sigurgeirsson *et al.*, 2016). Genes previously identified as differentially expressed between MSE and ESE were referred to as ‘known marker candidates’, including transcriptomic markers commonly used in endometrial testing (Díaz-Gimeno *et al.*, 2011; Enciso *et al.*, 2018), and others were recognised as ‘novel’.

In addition, the list of transcripts, significant in the unadjusted transcriptome and non-significant after deconvolution, were compared with gene expression profile of isolated epithelial and stromal cells, obtained from STRT RNA-seq (original data in Altmäe *et al.*, 2017). Genes, that exhibited no significant expression change on cell population level, were considered as genes whose expression change in whole tissue is derived from cell-type proportion fluctuations.

### Quantitative real-time PCR analysis

For technical validation of the results, reverse transcriptase quantitative PCR (RT-qPCR) was applied on selected endometrial marker candidates, using ten paired ESE and MSE total RNA samples from healthy volunteers different from these used in the RNA-seq study. The selection criteria for RT-qPCR validation were expression change |log_2_FC|≥1.5 in all groups and read count log_2_CPM≥4, based on our previous expertise.

1 µg of total RNA was reverse-transcribed using random hexamer primer (Thermo Scientific, USA) and cDNA was amplified and quantified using 7500 Fast Real-Time PCR System (Applied Biosystems, USA), using HOT FIREPol EvaGreen qPCR Mix Plus (Solis BioDyne, Estonia) and 250 nM of sense and antisense primers specific for nucleotide sequence of selected marker candidates or housekeeping genes, *TBP* and *SDHA*. Data were analysed using 7500 Software v2.0.5 (Applied Biosystems). Differences in gene expression levels were estimated using comparative Ct (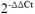) method (Livak and Schmittgen, 2001). Wilcoxon paired sample test was applied to estimate significance level. The oligonucleotide primers are presented in Supplementary Table S1.

## Results

### Differential expression analysis between mid-secretory and early secretory endometrium

In the present study, we used RNA-seq to analyse gene expression of endometrial samples. 70 individual cDNA libraries (35 ESE and 35 MSE samples) were sequenced and an average of 54 million read pairs were generated per library. The summary of RNA-seq results are provided in Supplementary Table S2. After filtering out low-quality reads from the datasets, an average of approximately 90% of the qualified read pairs across all groups were mapped to human genome version 19. Schematic overview of the analysis and results is presented in Figure 2.

**Figure 2.**
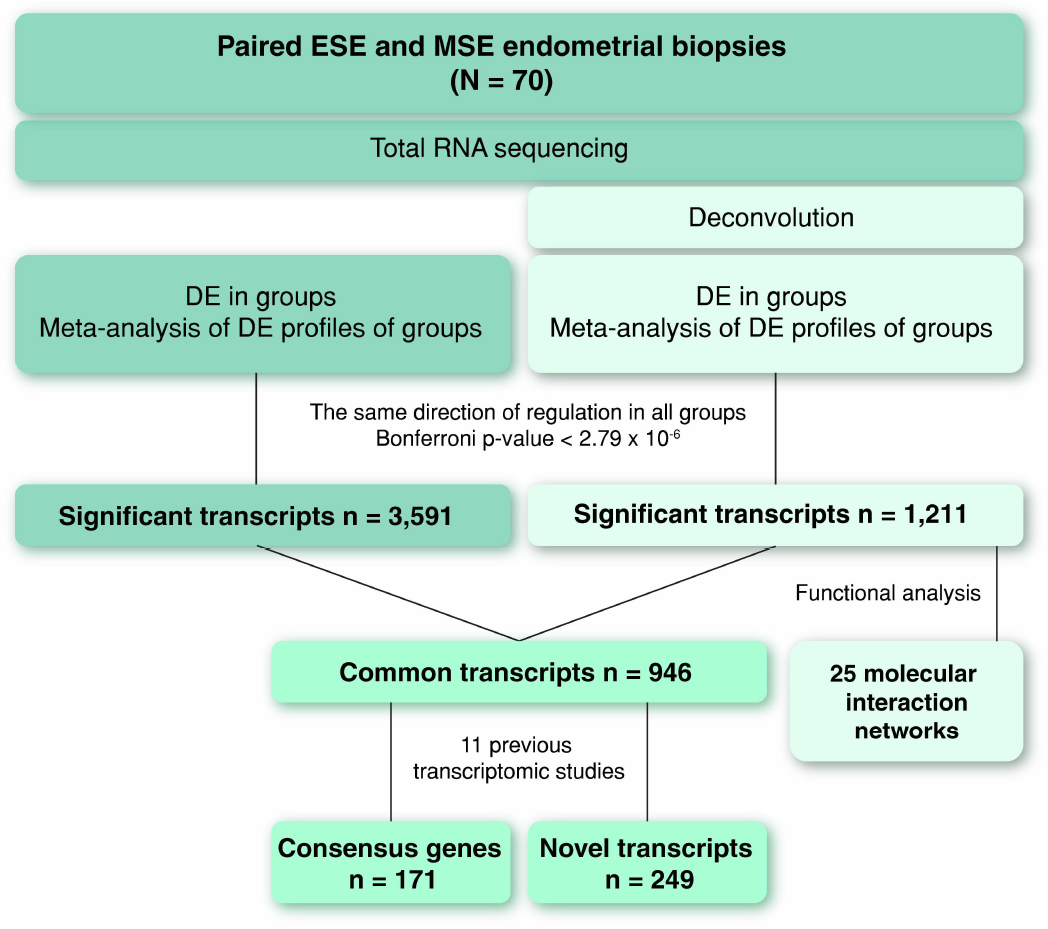
Overview of the endometrial RNA-seq data analysis pipeline and results. Sequenced endometrial total RNA samples were processed using two parallel approaches: i) differential expression analysis followed by meta-analysis of results from different groups and ii) deconvolution followed by DE and meta-analysis. Functional analysis was performed on all genes significant using deconvolution-based approach. Significant transcripts identified by two approaches were compared resulting in a set of common transcripts. ESE – early secretory endometrium; MSE – mid-secretory endometrium; N – number of analysed samples; n – numbers of DE transcripts identified during each step of the analysis; DE – differential expression; GO – Gene Ontology; Consensus genes – genes reported in 11 selected receptivity-related microarray and RNA-seq studies (please, see main text for the references).

The total number of expressed transcripts ranged from 21,890 to 30,576 per studied sample (Supplementary Table S2). DEGs within three groups were detected using paired sample design, followed by meta-analysis of all expressed genes with METAL software. This resulted in total of 3,591 genes (1,800 up-regulated and 1,791 down-regulated) that were identified as statistically significantly differentially expressed in MSE (Bonferroni p-value < 2.79 × 10^−6^) and showed the same direction in expression change in all three groups, as presented in the Supplementary Table S3. The average logarithmic expression fold change (Log_2_FC) of genes expressed in MSE ranged between 5.3 and 8.8 in both directions (Supplementary Table S3). Genes with the highest expression rate change were *PAEP, C4BPA, C2CD4A, CXCL14, GPX3, SLC15A1, MT1G, HAP1, MFSD4A* and *IRX3.* Most significantly differentially expressed genes were *SLC15A1, SLC1A1, SNX10, DKK1, GADD45A, AOX1, GPX3, DPP4, HEY2* and *MYOCD*.

### Expression profile deconvolution of endometrial tissue

To clarify if the observed changes in gene expression levels might actually stem from fluctuations in biopsy cellular composition, we used gene expression data for isolated endometrial epithelial and stromal cells to perform endometrial whole-tissue gene expression profile deconvolution. As a first step of the analysis, gene expression profiles of homogenous cell populations derived from ESE and MSE were compared with whole-tissue transcriptomes to estimate the relative proportions of these cellular subpopulations in the whole-tissue biopsy. This analysis revealed that the stromal fraction was larger in ESE biopsies, while in MSE biopsies the epithelial fraction was slightly larger, with considerable inter-individual variation (Figure 3A). On average, ESE samples consisted of 65% stromal cells, while the average stromal fraction in MSE samples was 46%. We observed a good correlation between gene-expression estimated epithelial cell proportions and endometrial cycle date according to histological evaluation (r=0.88 (95% CI 0.71-0.96), p=1.1 × 10^−6^), confirming that epithelial fraction becomes more dominant as the cycle progresses (Figure 3B).

**Figure 3.**
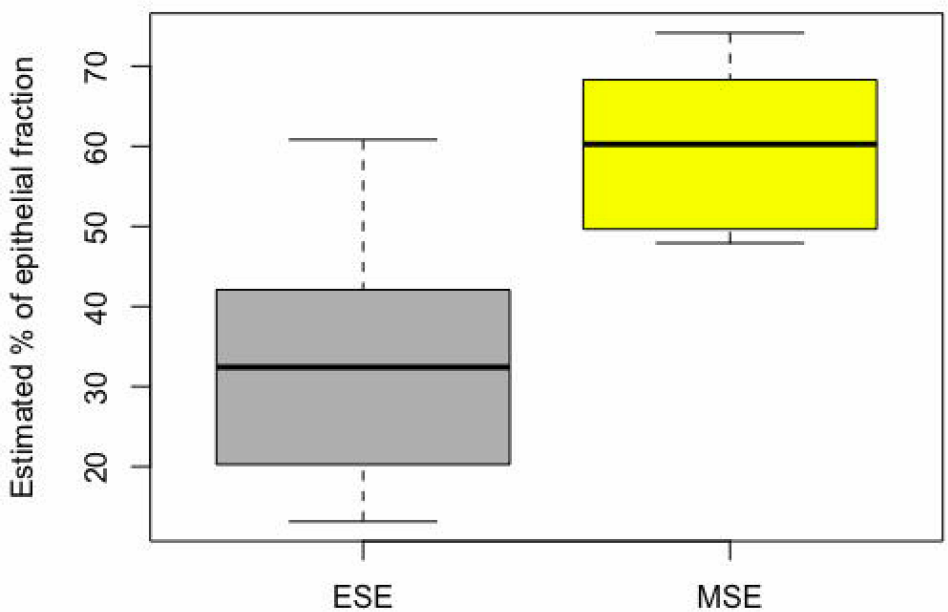
Tissue deconvolution results. A – Proportions of epithelial cells in endometrial biopsies estimated by computational deconvolution approach. B – Correlation between gene expression-estimated proportion of epithelial cells and menstrual cycle day predicted using Noyes’ criteria for endometrial histological dating. Black dots – ESE samples, red dots – MSE samples, according to urine LH test. C – Histological slide image of early secretory (ESE) and mid-secretory endometrial (MSE) samples, used for endometrial epithelial and stromal proportions’ counting. The samples were prepared from formalin-fixed paraffin-embedded endometrial tissue sections using standard hematoxylin and eosin staining protocol. E – epithelium and S – stroma. D – Comparison between epithelial cell proportions estimated using two different approaches, histological evaluation of epithelial fraction (dark grey) and proportions calculated based on cell-type specific gene expression patterns (light grey). In five out of six samples the proportions calculated by both methods were comparable.

As a second step, the epithelial and stromal cells’ proportions estimated by computational analysis of gene expression profiles were further validated by direct histological evaluation using endometrial tissue samples (Figure 3C). The histology results suggested that the proportion of epithelial cells in ESE is around 30%, increasing to approximately 50% by the time of WOI, and complies with the proportions calculated based on cell-type specific gene expression (35% and 54% for epithelial cells, respectively) (Figure 3D).

These proportions were then used as cofactors in differential expression analysis to account for differences in cellular composition. After meta-analysis of deconvoluted gene expression profiles from different groups, 1,211 (679 up- and 532 down-regulated) transcripts showed significant expression change in MSE endometrium, with Log_2_FC range between −5.3 and 7.9 (Supplementary Table S4). Genes with the highest expression rate change were *PAEP, HAP1, GPX3, CXCL14*, *C4BPA, SLC15A1, IRX3, MMP26, C2CD4A* and *SULT1C2P.* Most significantly differentially expressed genes were *SLC15A1, SLC1A1, SNX10, DKK1, GADD45A, AOX1, GPX3, DPP4, HEY2* and *MYOCD*.

As a result of RT-qPCR-based validation, *MYOCD, LINC01320, SLC8A1, TRPC4* and *GGATP1* showed significant expression rate change in MSE samples (Figure 4).

**Figure 4.**
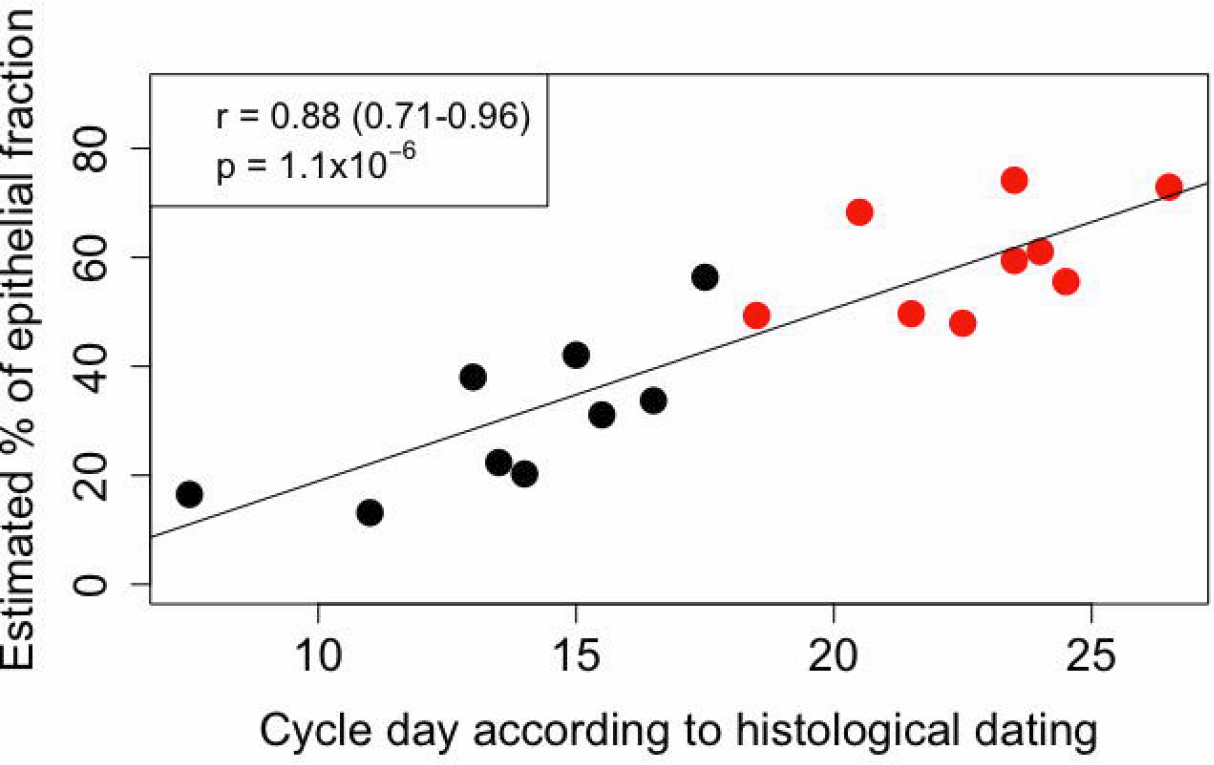
RT-qPCR validation of five marker candidates, using ten paired early secretory (ESE) and mid-secretory (MSE) endometrial samples. The dots connected with a solid line represent samples from the same patient. *Significant (paired Wilcoxon test p-value < 0.01).

### Transcripts differentially expressed in mid-secretory endometrium

All 1,211 transcripts that exhibited significant expression change in MSE after adjusting for differences in cellular composition were subject to functional annotation in order to investigate their role in endometrial maturation processes. DE genes with the highest significance level (n=90) and expression fold change (n=90) are depicted in Figure 5 (Supplementary Table S4).

**Figure 5.**
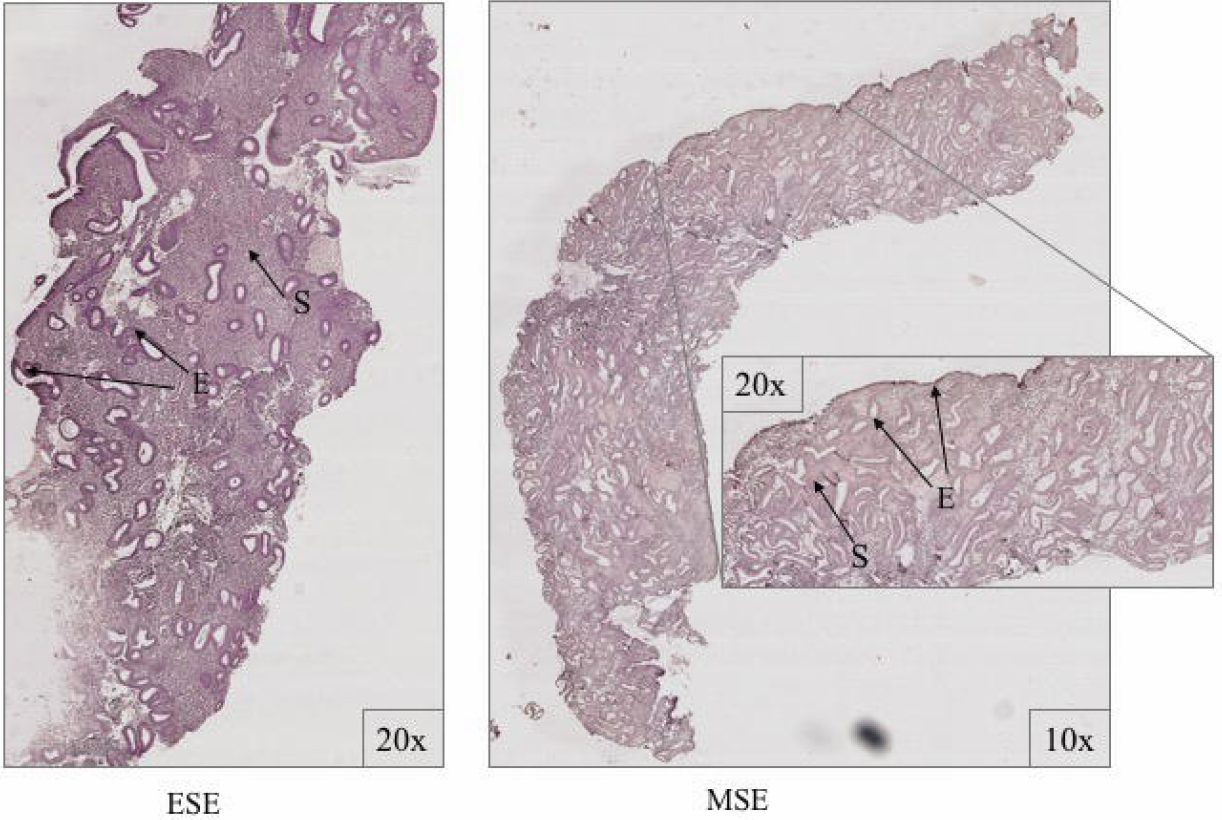
Differentially expressed genes (DEGs) between early secretory (ESE) and mid-secretory (MSE) endometrial samples. Scatterplot represents DEGs significant after deconvolution (Bonferroni p-value <2.79 × 10^−6^). Numbers represent chromosomes. Inner circle gene names contain top significant DEGs (red – up-regulated and blue – down-regulated in MSE, respectively), while outer circle contains DEGs with the highest expression rate fold change between ESE and MSE (pink – up-regulated and green – down-regulated in MSE, respectively).

Pathway mapping with IPA indicated that WOI DEGs are involved in many basic cellular functions and processes, and in particular cell fate, cellular movement, cell-to-cell signalling, cellular growth and development, inflammatory and vascular development processes (Supplementary Table S5). In total, IPA tool identified 25 Ingenuity Knowledge Database pathways significantly enriched in MSE transcriptome relative to ESE (Benjamini-Hochberg FDR< 0.05), highlighting the major regulative role of NFkB (Supplementary Figure S6 A), interferon a α (Supplementary Figure S6 B), steroid hormone activity (Supplementary Figure S6 C) and MAPK/ERK signalling (Supplementary Figure S6 D-F) in human endometrium. All molecular interaction networks with a score at least 21 (Krämer *et al.*, 2014) are presented in the Supplementary Materials (Supplementary Figure S6 A-K).

### Marker candidate genes for endometrial receptivity

When significant DEG lists before (3,591 transcripts) and after (1,211) deconvolution analysis were compared, 946 transcripts came out significant in both approaches (Figure 6, Supplementary Table S3 − Table S4) and therefore were considered as genes with the true expression change in MSE compared to ESE. 2,645 transcripts, which are almost 74% from all DE transcripts identified without deconvolution, were not considered significantly differentially expressed after adjustment for tissue cellular composition (Figure 6; Supplementary Table S3). Of these, genes with the highest expression change between MSE and ESE in whole tissue (Log_2_FC > 3), yet not differentially expressed after deconvolution, are presented in the Table 1. Moreover, 265 transcripts came out significant only after deconvolution was applied (Figure 6; Supplementary Table S4).

**Figure 6.**
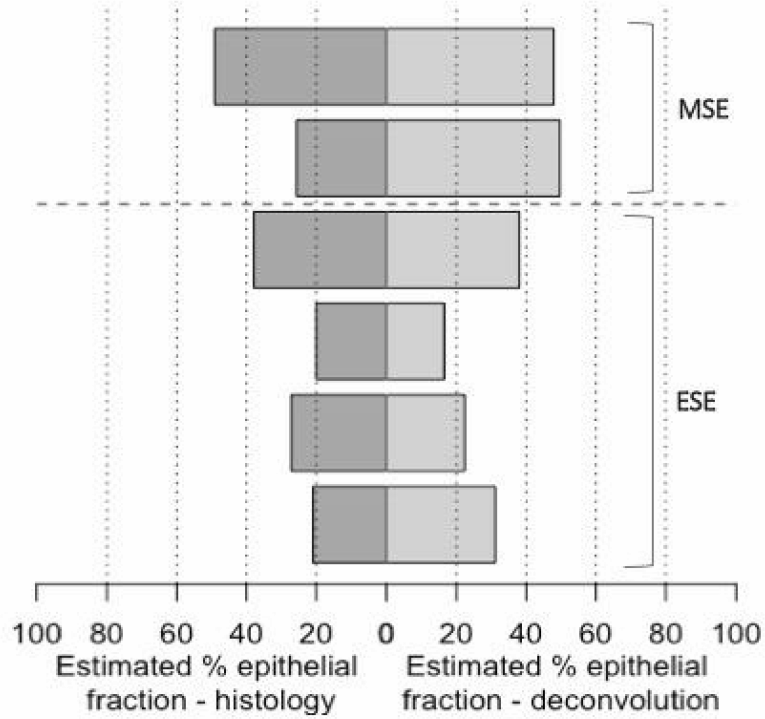
Numbers of significantly differentially expressed transcripts identified from meta-analysis of transcriptomic profiles of endometrial tissue samples from different groups. A – transcripts identified in whole endometrial tissue (before deconvolution, BD; n=3,591). B – transcripts identified in cellular heterogeneity-adjusted expression profiles (after deconvolution, AD; n=1,211). C – transcripts significant only in whole-tissue transcriptome and non-significant after deconvolution (blue, n=2,645); were considered genes, whose expression change in whole-tissue may be derived from variation in cell types’ proportions. D – Differentially expressed transcripts significant only after tissue deconvolution (pink, n=265); were considered genes whose expression change may be eclipsed in mid-secretory endometrium due to fluctuations in cell type proportions, leading to underestimation of molecular processes and pathways involved in endometrial maturation. E – transcripts significant in both approaches (purple, n=946) are considered potential WOI markers.

**Table 1.**
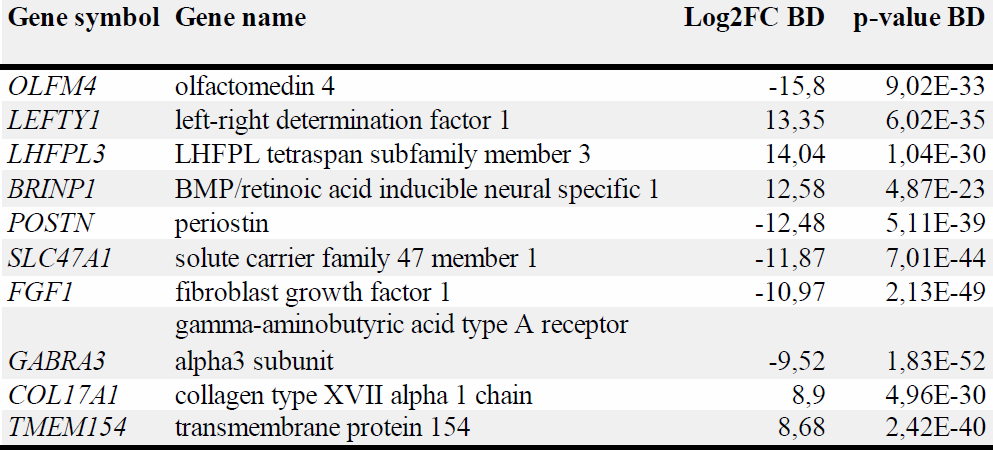
Top significantly differentially expressed genes in endometrial whole-tissue samples, that remained sub-significant after deconvolution analysis. Log2FC – average logarithmic expression fold change between mid-secretory and early secretory samples. P-value – Bonferroni p-value. BD – before deconvolution.

Further, we compared our results with those from eleven publications that have reported genes differentially expressed during WOI in healthy women, including nine microarray (Carson *et al.*, 2002; Kao *et al.*, 2002; Borthwick *et al.*, 2003; Riesewijk *et al.*, 2003; Mirkin *et al.*, 2005; Talbi *et al.*, 2006; Altmae *et al.*, 2010; Tseng *et al.*, 2010; Dìaz-Gimeno *et al.*, 2011) and two RNA-seq studies (Hu *et al.*, 2014; Sigurgeirsson *et al.*, 2016). Publications were selected based on the availability and quality of gene expression data sets. As a result, 171 transcripts overlapped among both microarray and RNA-seq studies (Supplementary Table S7 A), 526 transcripts were found in either microarray or RNA-seq study, while 249 transcripts were recognized as novel or previously not reported in endometrial WOI-related transcriptome (Supplementary Table S7 B). Top significant novel transcripts are presented in the Table 2.

**Table 2.**
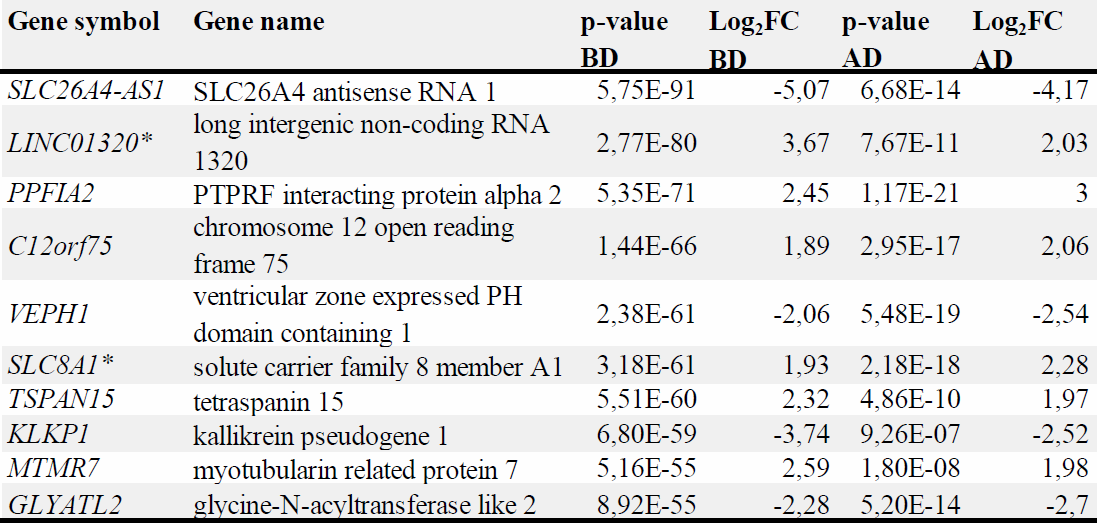
Top significant novel genes. Meta-analysis p-value using whole tissue expression profiling (Before Deconvolution, BD) and after adjusting for cellular composition (After Deconvolution, AD). Log2FC – average logarithmic expression fold change between mid-secretory and early secretory samples. * – RT-qPCR validated.

## Discussion

One of the main objectives of our study was to investigate how histological changes during menstrual cycle affect the detection of gene expression profiles of heterogeneous endometrial tissue. While the histological changes that distinguish MSE from ESE are well described (Noyes *et al.*, 1975; Crum *et al.*, 2003), to the best of our knowledge, we are the first to show how epithelial and stromal fractions are actually altered between pre-receptive and receptive state, by using computational deconvolution approach.

Since endometrial biopsies contain a range of cell types in varying proportions, due to both biological and technical reasons, some of the observed changes in gene expression levels might actually stem from differences in biopsy cellular composition, an issue that has not been addressed in previous endometrial transcriptome studies. As a substantial improvement compared to previous endometrial gene expression studies, we applied gene expression profile deconvolution to account for biopsy cellular composition that could artificially alter the observed expression profile of whole-tissue biopsy. Only 26% of the transcripts identified by meta-analysis without prior adjustment for cellular heterogeneity remained significant after deconvolution analysis was applied (Figure 6, Supplementary Table S3 and S4), while the remaining 74% rather reflect the variation in tissue cellular composition between the two compared time points. Among the latter category are much studied receptivity-associated genes *CLDN4, FOS, CCND2*, *AR, VC, CXCL13, PENK, OPRK1, KIF4A, POSTN, OLFM4, PRC1, KMO, AMIGO2, ECM1, RASSF2, KIF20A* and *LRP4* (Díaz-Gimeno *et al.*, 2011; Altmäe *et al.*, 2017). For example, olfactomedin 4 (*OLFM4*; Table 1) has been shown to be significantly down-regulated in MSE samples (Díaz-Gimeno *et al.*, 2011) but shows no significant expression change between MSE and ESE in either epithelial or stromal cells when evaluated separately (Dassen *et al.*, 2010; Kodithuwakku *et al.*, 2011).

Further, we described DE transcripts that were significantly differentially expressed only after deconvolution analysis was applied, whereas being non-significant before deconvolution (n=265, Figure 6). The expression change of these genes in whole-tissue biopsy may be eclipsed by fluctuations in cell type proportions in endometrial biopsy, potentially leading to underestimation of processes and pathways involved in endometrial maturation. Among these transcripts we found genes that have not been previously reported to be related to WOI. For example, cell migration-associated protein GLIPR2 and fibroblast growth factor 2 (FGF2) are believed to be involved in extracellular signal-regulated kinase (ERK) signalling (Paiva *et al.*, 2011; Huang *et al.*, 2013), one of the key pathways in endometrial gene expression regulation (Supplementary Table S5; Figure S6D) (Fluhr *et al.*, 2013). Elevated expression of vascular cell development-associated allograft inflammatory factor-1 (AIF-1) (Supplementary S6 K) was previously reported in human endometrium during luteal phase (Koshiba *et al.*, 2005) and was significantly up-regulated in our MSE samples. Inflammatory response-associated gap-junction alpha-1 (GJA1) (Supplementary S6 D), also found significantly DE only after deconvolution analysis, is differentially expressed in stromal cells during MSE, promotes stromal cell differentiation and maintenance of decidualized endometrial phenotype (Yu *et al.*, 2011, 2014). Although, to the best of our knowledge, these genes have not yet been reported in human MSE transcriptome studies before, while their aberrant expression has been shown in endometrial studies focusing on implantation failure, insufficient endometrial growth and antiprogestin effect (Catalano *et al.*, 2007; Savaris *et al.*, 2008; Tapia *et al.*, 2008; Maekawa *et al.*, 2017), supporting their involvement in endometrial processes. It is important to outline that in our study these genes did not show significant expression change in whole-tissue samples, suggesting that dynamic epithelial and stromal cell proportions may also be the reason they have not been reported before in whole-tissue transcriptomic studies.

The information about the endometrial receptivity-associated gene expression keeps improving. Thousands of genes have been suggested to take part in the processes of endometrial maturation, with respect to their WOI-marker potential. Previous findings summarized in two latest publications have outlined that WOI could be characterized by a relatively small number of differentially expressed genes (Altmäe *et al.*, 2017; Enciso *et al.*, 2018). In the present study, we also report a set of 1,211 transcripts differentially expressed between MSE and ESE after deconvolution (Supplementary Table S4). Figure 5 summarizes the set of transcripts, significantly differentially expressed in MSE after endometrial gene expression profile was adjusted for epithelial and stromal fractions, by extracting the transcripts with the highest expression rate change (outer gene circle, n=90) and significance level (inner gene circle, n=90). These top genes also belonged to 946 potential endometrial receptivity markers that exhibited significant expression change in whole-tissue transcriptome analysis. Moreover, 31 genes (*C4BPA, CD55, DPP4, AOX1, LMCD1, PLCH1, FRAS1, BMPR1B, SH3RF2, GPX3, SOD2, CYP2A5, SNX10, PRR15, SLC1A1, PAEP, DKK1, HABP2, DLG2, NNMT, TRPC4, SLC15A1, NPAS3, FGF7, C2CD4A, IRX3, MYOCD, PAK5, POM121L9, MAOA* and *GRIA3*) were located in both circles, that brings up their great biomarker potential. Twelve of these genes, *PAEP, GPX3, C4BPA, DPP4, DKK1, CD55, NNMT, MAOA, HABP2, SOD2, AOX1* and *SLC1A1*, are already applied endometrial receptivity markers (Díaz-Gimeno *et al.*, 2011; Altmäe *et al.*, 2017; Enciso *et al.*, 2018), while others have been previously reported in endometrial receptivity-related transcriptome studies (Altmäe *et al.*, 2012; Hu *et al.*, 2014; Sigurgeirsson *et al.*, 2016; Chen *et al.*, 2017a). Still, many of the genes identified in our study (Supplementary Table S4) have not been reported in MSE transcriptome studies before. For example, solute carrier family member, *SLC8A1*, one of the top differentially expressed genes, is recruited in ERK1/2 activation and can modulate immunoglobulin-mediated immune response through ERK1/2 or through interaction with major histocompatibility complex (MHC) class I proteins (Supplementary Table S5, Figure S6 D and Table S7 B). Up-regulated in stromal cells cathepsin B (CTSB) suggests its non-proteolytic activity (Sigloch *et al.*, 2016) leading to promoted collagen type II expression and MAPK/ERK activation (Supplementary Figure S6 F and Table S7 B).

Our results also support previously discussed important role of immune response in pre- and peri-implantation period (Altmäe *et al.*, 2010; Gnainsky *et al.*, 2014; Haller-Kikkatalo *et al.*, 2014; Altmäe *et al.*, 2017). For example, type I interferon-activated proteins caspase 1 (CASP1), tumour suppressor protein SAMD9L and phospholipid scramblase 1 (PLSCR1) induce innate immune response and promote endometrial antiviral status (Catalano *et al.*, 2007; Boon *et al.*, 2014; Wang *et al.*, 2016) (Supplementary Figure S6 B). We also noted that the expression change of these three genes in MSE before deconvolution was under 2-fold (Supplementary Table S3), which may be the reason why these genes have never been reported in studies using at least 2-fold expression change threshold. The 171 DE genes, overlapping between our results and previously published RNA-seq and array studies, can be considered as genes with the highest endometrial receptivity marker potential, while novel marker candidates reported in our study need further investigation in order to understand how they are involved in endometrial maturation (Supplementary Table S7 A and B).

With the implementation of transcriptomic tools in endometrial analysis, it has become clear that not only protein coding genes are important for endometrial receptivity. Long inter-genic noncoding RNAs (lincRNAs) have significant regulative potential on transcriptional and translational level through direct interactions with different types of RNAs and proteins (Kung *et al.*, 2013; Groen *et al.*, 2014). Recently, Sigurgeirsson and colleagues showed the potential role of lincRNAs in endometrial receptivity (Sigurgeirsson *et al.*, 2016). We report eight lincRNAs significantly up- or down-regulated in MSE (Table 3). Our results demonstrate that *LINC01320* in particular has a strong effect in all studied groups (Table 2, Supplementary S3-S4), confirmed by RT-qPCR (Figure 4). Previously, differential expression of *LINC01320* was reported in endometrial and ovarian cancer cells (Chen *et al.*, 2017a), although, to the best of our knowledge, it has never been associated with endometrial receptivity before.

**Table 3.**
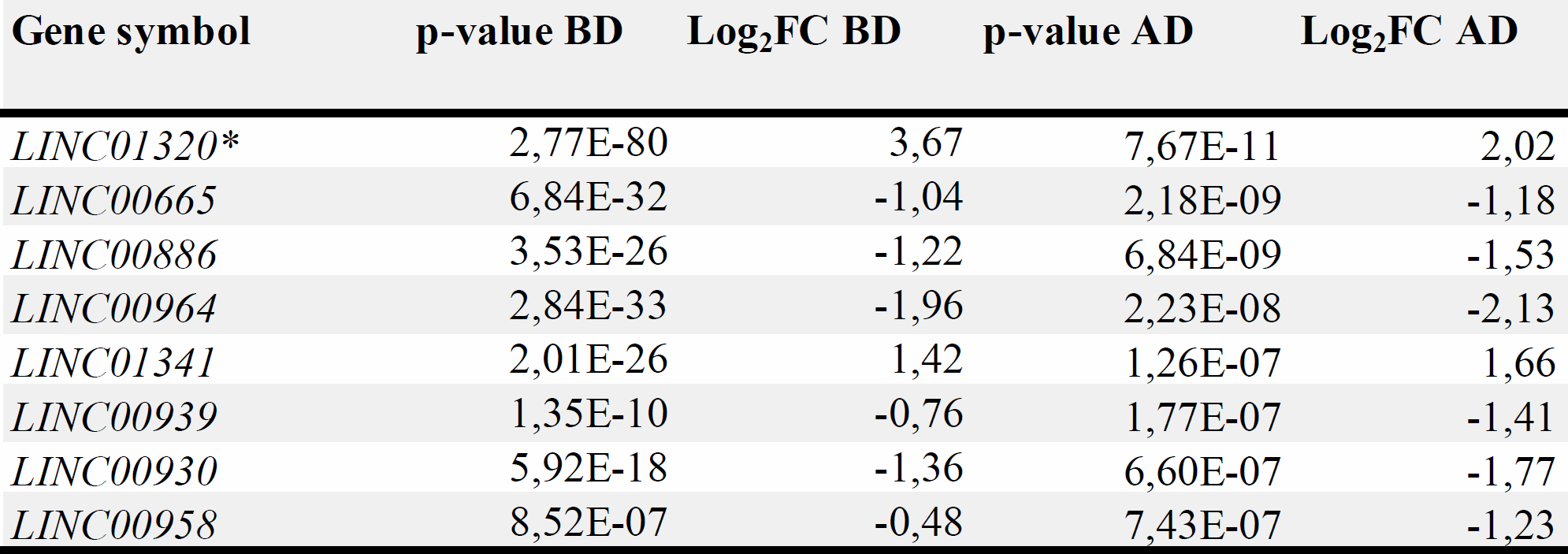
Long intergenic non-coding RNAs, significantly differentially expressed in human endometrium during WOI. Log2FC – average logarithmic expression fold change between mid-secretory and early secretory samples. P-value – Bonferroni p-value. BD -before deconvolution. AD – after deconvolution. * – RT-qPCR validated.

Certain limitations of our study also need to be acknowledged. For gene expression profile deconvolution, only stromal and epithelial cell fractions were considered, whereas other cells, such as endothelial and immune cells, also contribute to the gene expression profile. Second, using actual counts for cellular subpopulations obtained by FACS as covariates in differential expression analysis is likely to provide more accurate expression change estimates than computational deconvolution applied in this study. Nonetheless, computational deconvolution is becoming increasingly popular (Reinartz *et al.*, 2016; Scott *et al.*, 2016; Yu and He, 2017), and has been shown to have accuracy similar to methods using actual cell counts (Newman *et al.*, 2015). As a marked methodological improvement, we address the transcriptomic changes in human endometrium using meta-analysis of samples from different populations, which not only helps to further eliminate population-specific and batch effects, but also to gain the statistical power required. As a result, we suggest a set of robust endometrial receptivity markers that reflect actual expression changes that take place in endometrial tissue, irrespective of its cellular composition. However, novel potential endometrial transcriptomic markers identified in our study, such as *LINC01320*, need further investigation to elucidate their role in endometrial functioning.

In conclusion, our results show how the proportions of endometrial epithelial and stromal cells change in endometrial biopsies during transition from pre-receptive to receptive state and demonstrate how these changes affect gene expression profile of biopsied tissue. Overall, our results highlight that biopsy cellular composition must be taken into consideration in endometrial ‘omics’ studies where whole tissue samples are involved.

### Acknowledgements

We thank Dr. Peeter Karits and Dr. Elle Talving as well as the personnel at Nova Vita and BioEximi fertility clinics for their involvement in sample collection, Katrin Kepp for the recruitment of healthy women and Dr. Keiu Kask for the help with histological imaging. We are also thankful to the personnel at IVI clinic Valencia for their involvement in sample collection, and to Dr. Pilar Alamá for the recruitment of healthy volunteers. We express our gratitude to all study participants from both countries.

### Funding

This study was funded by Estonian Ministry of Education and Research (grant IUT34-16); Enterprise Estonia (grant EU48695); the EU-FP7 Eurostars program (grant NOTED, EU41564); the EU-FP7 Marie Curie Industry-Academia Partnerships and Pathways (IAPP, grant SARM, EU324509); Horizon 2020 innovation program (WIDENLIFE, EU692065) and MSCA-RISE-2015 project MOMENDO (grant no 691058). FV was supported by the Miguel Servet Program Type I of Instituto de Salud Carlos III (CP13/00038). Authors confirm no competing interests.

## Supplementary material

S1 Primers for RT-qPCR

S2 RNA-seq results

S3 DE transcripts before deconvolution

S4 DE transcripts after deconvolution

S5 Functional analysis

S6 IPA Networks

S7 Consensus and novel transcripts

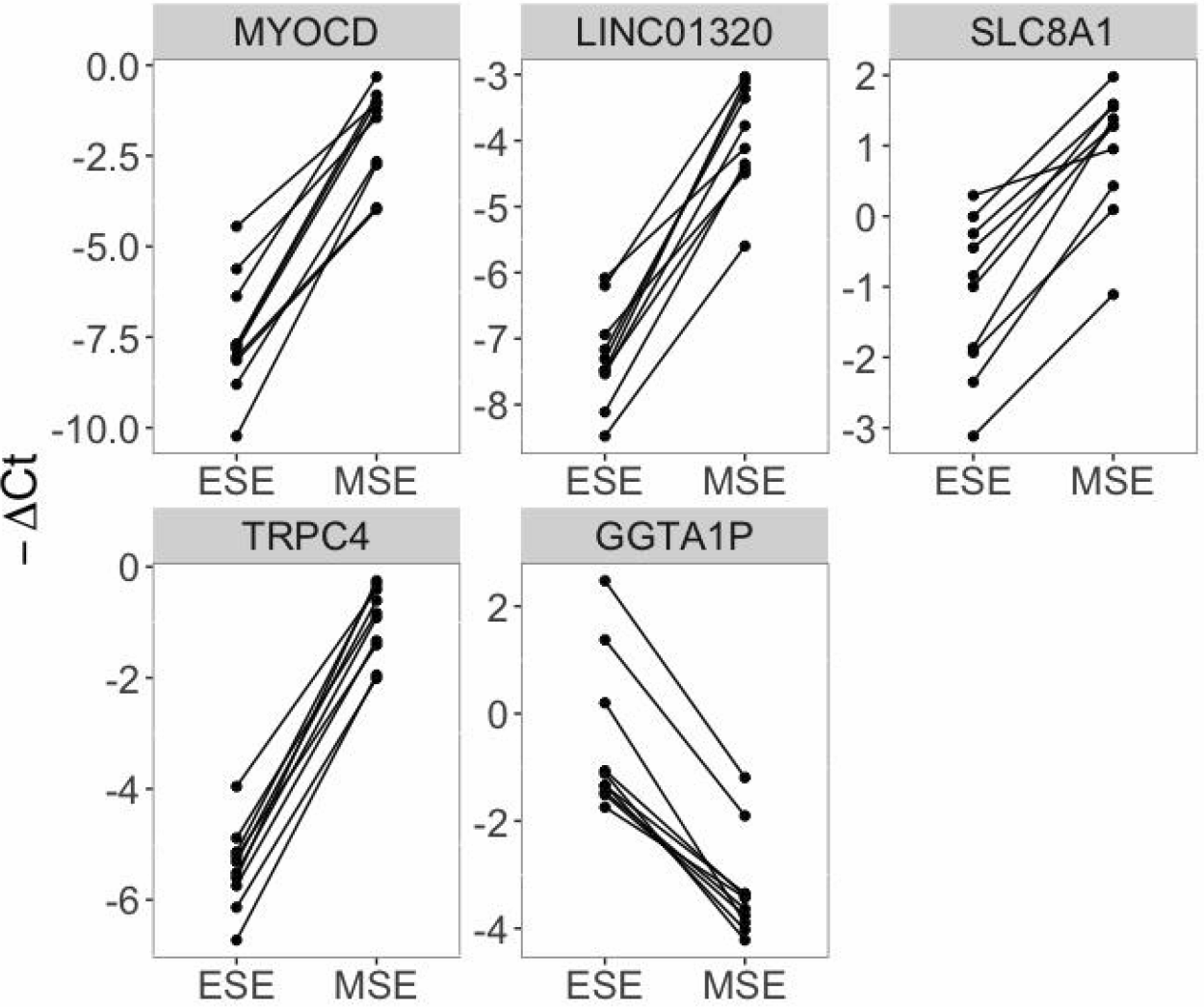

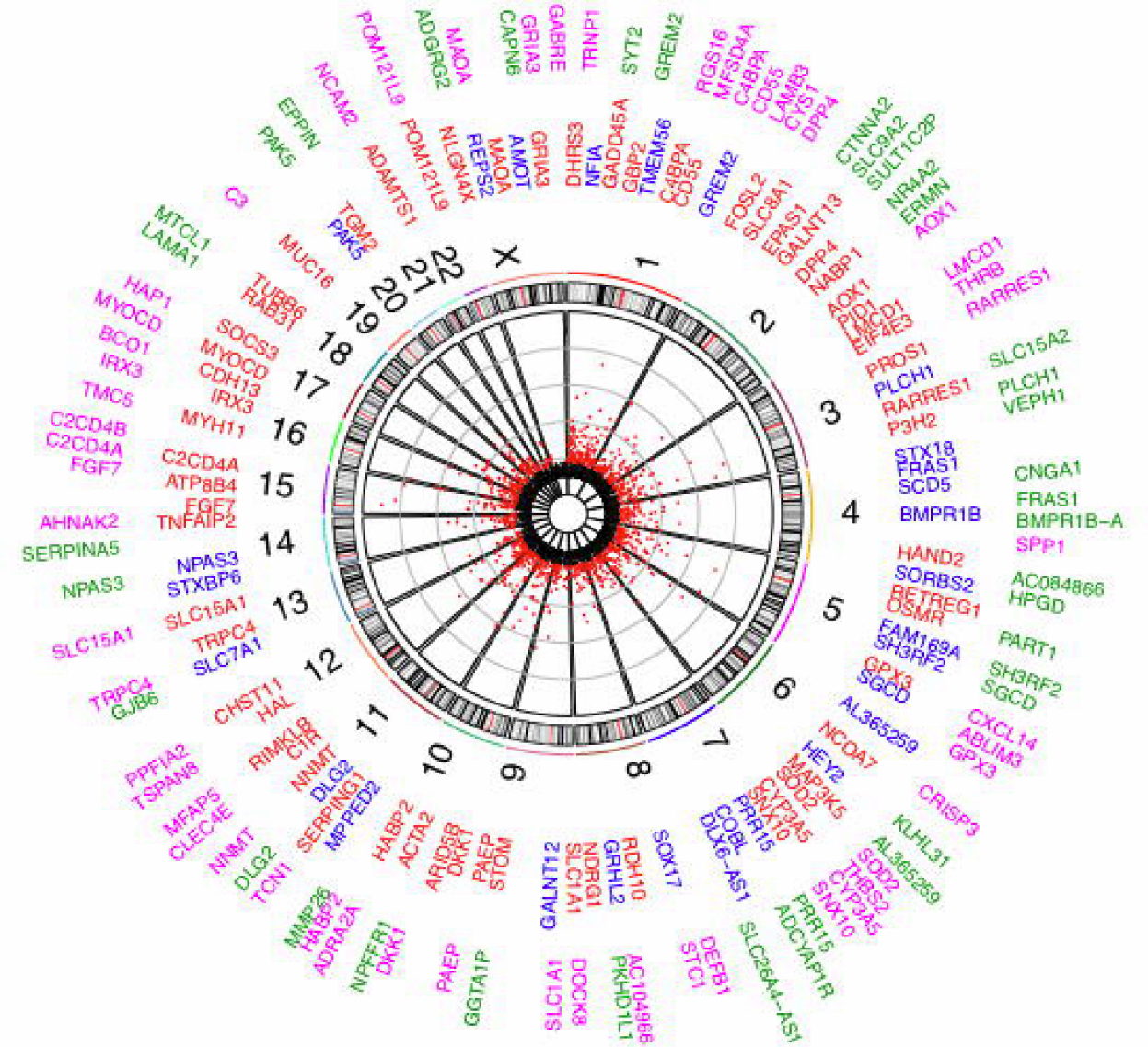

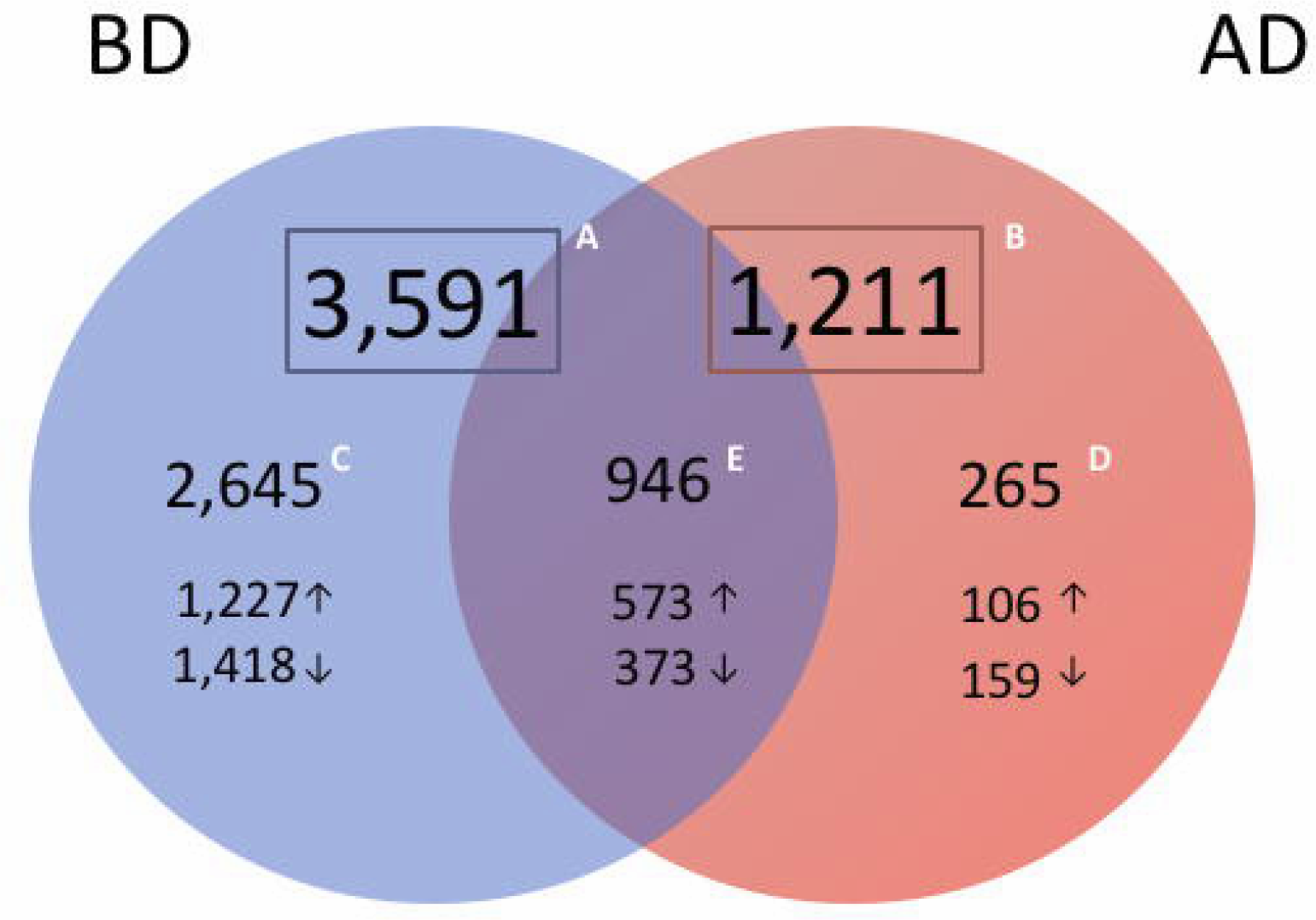

